# Lacking conservation genomics in the giant Galápagos tortoise

**DOI:** 10.1101/101980

**Authors:** Etienne Loire, Nicolas Galtier

## Abstract

This preprint has been reviewed and recommended by Peer Community In Evolutionary Biology (http://dx.doi.org/10.24072/pci.evolbiol.100031).

Conservation policy in the giant Galápagos tortoise, an iconic endangered animal, has been assisted by genetic markers for ∼15 years: a dozen loci have been used to delineate thirteen (sub)species, between which hybridization is prevented. Here, comparative reanalysis of a previously published NGS data set reveals a conflict with traditional markers. Genetic diversity and population substructure in the giant Galápagos tortoise are found to be particularly low, questioning the genetic relevance of current conservation practices. Further examination of giant Galapagos tortoise population genomics is critically needed.

---

Biological conservation implies collaboration between scientists, biodiversity managers and policy makers. We can only applaud when scientific progress influences policies and practices in a way that is beneficial to nature, and ultimately to humans. A recent Comment in Nature (Nichols 2015) highlights a spectacular conservation initiative geared towards an iconic endangered species – the giant Galápagos tortoise.

The Galápagos Islands host a number of fascinating endemic animals, such as Darwin’s finches, Galápagos iguanas and Galápagos penguins and their preservation is clearly a major issue. These Islands are a paramount conservation priority. Remarkable efforts have been made by the Charles Darwin Research Station and the Galápagos National Park Service to protect indigenous species while eradicating invasive ones (e.g. Marquez et al. 2013). The Galapagos tortoise, *Chelonoidis nigra*, is probably the most emblematic of all species of the archipelago. These large tortoises have been heavily impacted by human activities over the centuries. Since the mid-1960s they have been the focus of a specific long-term conservation program, which has been fully successful in terms of population density recovery (e.g. Gibbs et al. 2014, Aguilera et al. 2015). The program mainly involved bringing eggs or adults from natural sites to rearing/breeding centers, and releasing in the field juveniles of sufficient size, thus escaping predation by rats (Cayot 2008).

Since the 2000’s, population genetics research has been instrumental in the management plan of Galápagos tortoises. Thirteen giant Galápagos tortoise species (Poulakakis et al. 2015) have been described, or confirmed, using genetic data – specifically, mitochondrial DNA and 9-12 microsatellite loci (Caccone et al. 2002, Ciofi et al. 2002). Based on this information, population densities have been monitored in a way that aims at preserving, or even restoring, the integrity of the distinct gene pools (e.g. Milinkovitch et al. 2004, Garrick et al. 2012, Edwards et al. 2014). Milinkovitch et al. (2007), for instance, identified one “hybrid” individual among the thousands that had been repatriated to the Española island after captive breeding. This was removed from the field. Garrick et al. (2012) and Edwards et al. (2013) discuss the “conservation value” of individuals based on their genotypes. The most recent implementation of this strategy is an attempt to recreate the genome of the famous Lonesome George tortoise, the last member of the *C*. (*nigra*) *abingdoni* species, which died in 2012. The plan is to cross individuals from another species that carry *abingdoni* alleles – hence the recent transfer by helicopter and boat of 32 tortoises from their current habitat to the island of Santa Cruz (Nichols 2015).

Some problems, however, still hamper the task of defining *C. nigra* taxonomic and conservation units. For instance, mitochondrial DNA and microsatellite data do not fully agree (Poulakakis et al. 2012). Published levels of mitochondrial coding sequence divergence between named species of giant Galápagos tortoises (<1%, Caccone et al. 2004) are not higher than typical amounts of within- species mitochondrial polymorphism in reptiles (Bazin et al. 2006). Importantly, individuals carrying so-called ‘hybrid’ genotypes or unexpected mitochondrial haplotypes, given their geographic origin, are common in the field (e.g. Garrick et al. 2012). Because they are interpreted as being the result of non-natural causes (Russello et al. 2007, Poulakakis et al. 2008), such individuals have been removed from published analyses (Garrick et al. 2014, Poulakakis et al. 2015), thus artificially inflating measured levels of genetic differentiation between entities. In spite of this bias, estimated FST as low as 0.1 were considered sufficient to delineate species (Poulakakis et al. 2015), in absence of evidence for hybrid depression or local adaptation. Indeed, very little is known regarding the link between genetic variation, morphological variation and fitness in *C. nigra*. The conspicuous “saddleback” morphotype, for instance, is observed in several unrelated species that carry various combinations of mtDNA haplotypes and microsatellite alleles (e.g. Poulakakis et al 2012), and the heritability of this trait is so far unknown (Chiari et al. 2009). Whether gene flow, if any, is recent and human mediated, or ancient and “natural”, is critical with respect to conservation policy in *C. nigra*.

Four years ago, we published the blood transcriptome of five giant Galápagos tortoises which, based on mtDNA, were assigned to three distinct species – *becki*, *vandenburghi*, and *porteri* (Loire et al.2013). To our surprise, our analysis of ∼1000 coding loci did not reveal any evidence of genetic structure in this sample, i.e. the homozygosity of the five individuals was not higher than expected under the hypothesis that they belonged to a single panmictic population. The total genetic variation detected in the sample was also quite low – heterozygosity was around 0.1%, consistent with earlier findings (Caccone et al. 2004).

Here we looked at the situation from a comparative standpoint and plotted the inbreeding coefficient F as a function of log-transformed synonymous diversity, πS, in a dataset of 53 species of animals for which the transcriptome of at least four individuals sampled from different populations was available (Perry et al. 2014, Romiguier et al. 2014, Table S1). The synonymous diversity *π*_S_ measures heterozygosity at (presumably neutral) third codon positions; *π*_S_ is an unbiased estimate of the *θ*=4.N_e_.*μ* parameter, where N_e_ is the effective population size and *μ* is the per-base, per-generation mutation rate. Inbreeding coefficient F measures the departure from Hardy-Weinberg equilibrium by comparing the expected and observed numbers of heterozygous genotypes, here summed across individuals and loci. F is expected to equal zero in absence of population structure, and to take positive values when individuals are more homozygous, on average, than expected under the assumption of panmixy. Data on all species in this dataset were analysed via the same procedure (Gayral et al. 2013, Ballenghien et al. 2017), thus ensuring optimal comparability between data points. Figure 1 indicates that *C. nigra* harbours lower genetic polymorphism and population structuring than most vertebrates or other animal species, which casts serious doubts on the existence of several differentiated gene pools in this taxon. Interestingly, the data point closest to*C. nigra* in this figure is *Homo sapiens* – a species in which very few people would argue the need to define (sub)species.

**Figure 1.**
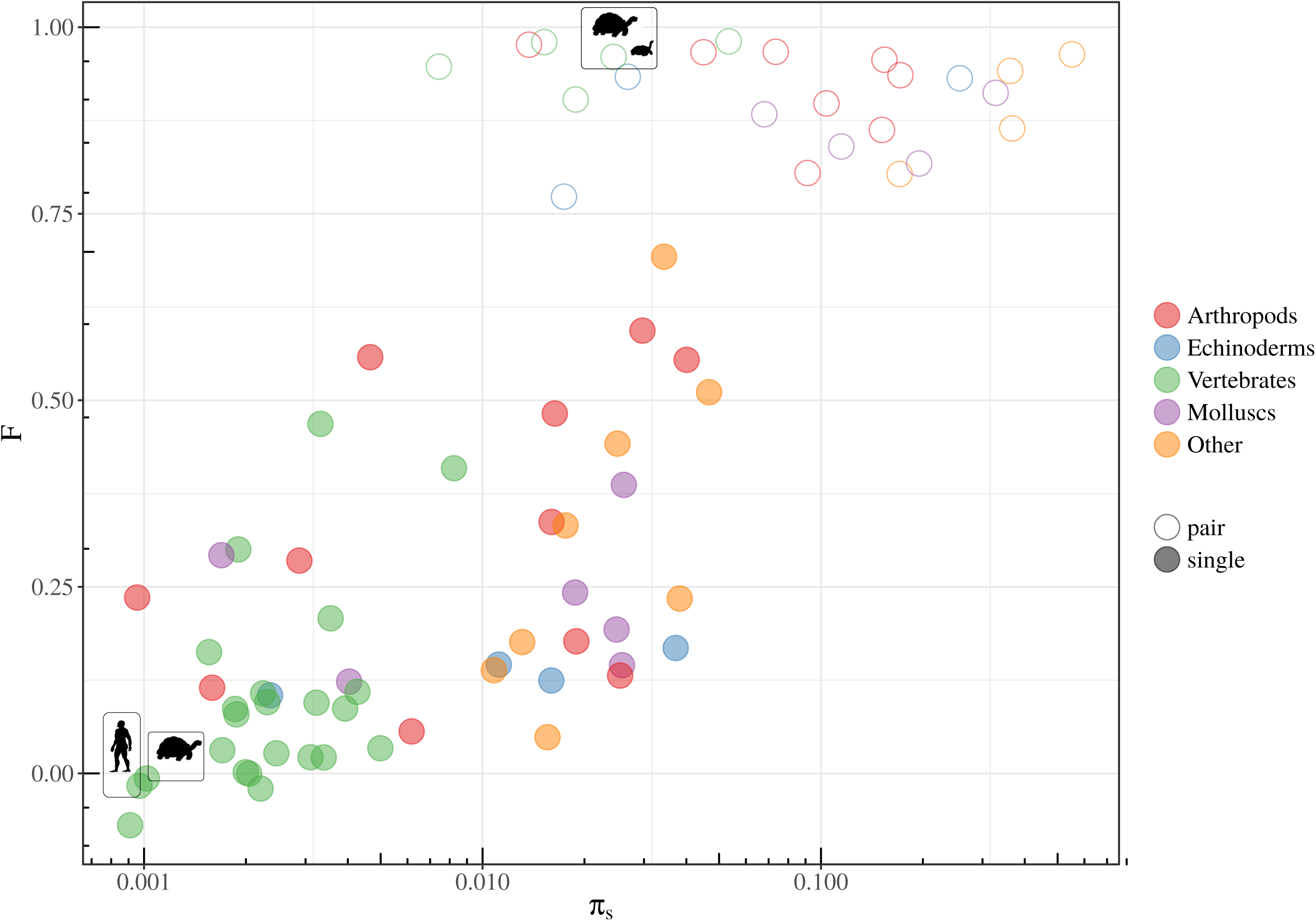
Genetic diversity and population substructure in 53 species and 25 pairs of congeneric species of animals. Each filled circle corresponds to a single species of animals in which the transcriptome of at least four individuals was sequenced (Perry et al. 2012, Romiguier et al. 2014). Harvest ant *Messor barbarus* was excluded due to its highly specific mating system (Romiguier et al. 2017). Each open circle corresponds to a pair of congeneric species of animals in which the transcriptome of at least five individuals was sequenced (Galtier 2016). Genotypes were called using the improved, contamination-aware algorithm of Ballenghien et al. (2017), parameter γ being set to 0.2. X-axis (log scale) reflects the amount of neutral genetic variation within the sample. This is measured by the synonymous-site heterozygosity, *π*S, for single species, and by the proportion of fixed synonymous differences for species pairs. Y-axis reflects the detected amount of population substructure, measured by the F statistics. The positions in the graph of *C. nigra*, *Homo sapiens* (single species) and of the *C. nigra/Chelonoidis carbonaria* pair are indicated by thumbnails.

The above analysis is based on small samples – from five to eleven individuals per species – which in most cases precluded definition of sub-populations and calculation of the F_ST_ statistics – but note that a significantly positive FST, if any, should imply a significantly positive F, and that F in this analysis was calculated using the method of Weir and Cockerham (1984), which accounts for small sample size. Our analysis might be affected by, *e.g*., sequencing errors, genotyping errors, hidden paralogy, cross-contamination or other experimental artefacts (Ballenghien et al. 2017). To control for possible sources of noise and confirm that our approach has the power to identify strong genetic structure when it exists, we similarly analysed samples made of individuals from two congeneric species (Galtier 2016, Table S1). Distinctively high values of F were obtained with the two-species samples, compared to the single-species ones (Figure 1, open circles), casting further doubts on the hypothesis that our *C. nigra* sample comprises individuals from more than one species.

It was suggested from mtDNA and microsatellite data that some *C. nigra* individuals are of “hybrid origin”, i.e., F1 from “purebred” parents or backcrosses (e.g. Russello et al. 2010, Garrick et al. 2014). Among the four individuals of our sample for which microsatellite data are available, two were consistently assigned to the same population by the three methods used by Russello et al. (2010), whereas two obtained conflicting assignments. Russello et al. (2010) analyzed 156 individuals of unknown geographic origin. Re-analysing their Table S1, we found that 109 of these (70%) were assigned to two or three distinct populations by the three methods they used. Our sample, therefore, is not particularly enriched in individuals of “hybrid origin”. Still, to control for a possible impact of hybrids in our analysis, we calculated the maximal F across the ten possible pairs of individuals from our sample. If any two individuals from our sample belonged to distinct species, this statistics should take a high value. We rather found that the maximal pairwise F equals a low 0.12, indicating that the absence of homozygote excess in *C. nigra* (Figure 1) is not explained by the presence of a few “hybrids” in our sample, but rather reflects an overall lack of genetic differentiation.

The sample of Loire et al. (2013) is small in terms of the number of individuals and obviously insufficient to provide a definitive answer regarding population genetic structure in *C. nigra*. Our sample, however, is large with regard to the number of loci. Distinct loci independently sample the coalescent process, and therefore provide small but many pieces of information on population history. The strength of even small-sized samples to reconstruct population processes, provided that a large number of loci is available, has been demonstrated in a number of recent publications (e.g. Der Sarkissian et al. 2015, Abascal et al. 2016). The popular PSMC method (Li and Durbin 2011), for instance, estimates former fluctuations in effective population size based on a single diploid genome. Roux et al. (2016) specifically demonstrated the potential of the Romiguier et al. (2014) data set to inform on species boundary and the history of gene flow between populations.

Our results, if confirmed, are highly relevant for the conservation of Galápagos tortoises. They could imply that the current practice whereby only individuals from the same “species” are crossed is not only pointless – in the absence of truly differentiated gene pools – but also potentially harmful to the species due to the risk of inbreeding depression. The risk is particularly high when large numbers of closely related individuals are simultaneously released in a closed area (Milinkovitch et al. 2004). In the same vein, physical removal from the field of naturally occurring ‘hybrids’ (Milinkovitch et al. 2007) and sterilization of individuals to avoid undesired crosses (Hunter et al.2013) might well actively participate in the decline in the mean fitness of local populations (Frankham et al. 2014).

Of note, even the most recent publications on this issue, including the description of a new species (Poulakakis et al. 2015) have focused on the same limited set of markers developed some 15 years ago. We suggest that, nowadays in the genomic era, the conservation policy regarding Galápagos tortoises should no longer be based entirely on just a dozen loci, given the demonstrated potential of genome-wide data to enhance population genetics research in other groups of threatened animals (e.g. Liu et al. 2014, Moore et al. 2014, Der Sarkissian et al. 2015, Dobrynin et al. 2015, Lamichaney et al. 2015, Abascal et al. 2016). Knowing the importance of inbreeding depression as a threat to conservation of endangered species (e.g. Allendorf et al. 2010, Walling et al. 2011, Frankham 2015) we suggest that forced inbreeding and sterilization interventions in Galápagos tortoises be delayed until the genetic architecture of the species is clarified via the analysis of genome-wide data in a sufficiently large sample, thus resolving the discrepancy between existing datasets. There is some controversy and subjectivity in the definition of relevant conservation units and strategies (Edmands 2007, Funk et al. 2012, Shaffer et al. 2015, Taylor et al. 2017). Giant Galapagos tortoises are maximally protected throughout the archipelago. The definition of conservation units in *C. nigra*, therefore, is not relevant to the preservation of local populations. Rather, genetic data have been used to make decisions on which individuals should be crossed with which, and where they should be located. This is only meaningful if the distinct entities are sufficiently isolated, genetically and geographically, such that gene flow is naturally low. This is why species boundaries matter: current conservation goals in *C. nigra* are dependent on the validity of the existing population model and taxonomy.

It is likely that human intervention during the last centuries has not only dramatically affected *C. nigra* population density, but also favoured migration of individuals across islands. It is also quite likely, based on existing data, that gene flow between islands and/or local extinction/recolonization was substantial before human intervention, preventing speciation in *C. nigra* – otherwise hybrids would not be so common, total genetic diversity/divergence would be higher, population substructure would be stronger and more obviously correlated to morphology. In other words, *C. nigra* might well be better described as a perturbed metapopulation than a collection of endangered species. If this was true then the goals of resurrecting ancestral “lineages” and preventing crosses between genetic entities would just be nonsense. Only a genome-wide analysis has the power to discriminate between ancient and recent gene flow, clarify the link between genotype and phenotype, and inform conservation policies. Current management aims at preserving/restoring particular combinations of alleles at 12 specific loci; relevance to phenotypes, fitness, population history and species barriers in *C. nigra* is anything but clear.

